# *TAF1*-gene editing impairs Purkinje cell morphology and function

**DOI:** 10.1101/545491

**Authors:** Udaiyappan Janakiraman, Jie Yu, Aubin Moutal, Shelby N. Batchelor, Anandhan Annadurai, Rajesh Khanna, Mark A. Nelson

**Affiliations:** Departments of Pathology, University of Arizona College of Medicine and College of Pharmacy, Tucson, AZ, USA; Departments of Pharmacology, University of Arizona College of Medicine and College of Pharmacy, Tucson, AZ, USA; Department of Pharmacology and Toxicology, University of Arizona College of Medicine and College of Pharmacy, Tucson, AZ, USA; The Center for Innovation in Brain Sciences, The University of Arizona Health Sciences, Tucson, Arizona; The BIO5 Institute, University of Arizona

**Keywords:** TAF1, Intellectual disability syndrome, CRISPR/Cas system, X-linked disorder, cerebellum

## Abstract

TAF1 intellectual disability syndrome is an X-linked disorder caused by loss-of-function mutations in the TAF1 gene. How these mutations cause dysmorphology, hypotonia, intellectual and motor defects is unknown. Mouse models which have embryonically targeted TAF1 have failed, possibly due to TAF1 being essential for viability, preferentially expressed in early brain development, and intolerant of mutation. Novel animal models are valuable tools for understanding neuronal pathology. Here, we report the development and characterization of a novel animal model for TAF1 ID syndrome in which the *TAF1* gene is deleted in embryonic rats using clustered regularly interspaced short palindromic repeats (CRISPR) associated protein 9 (Cas9) technology and somatic brain transgenesis mediated by lentiviral transduction. Rat pups, post-natal day 3, were subjected to intracerebroventricular (ICV) injection of either gRNA-control or gRNA-TAF1 vectors. Rats were subjected to a battery of behavioral tests followed by histopathological analyses of brains at post-natal day 14 and day 35. *TAF1*-edited rats exhibited behavioral deficits at both the neonatal and juvenile stages of development. Deletion of TAF1 lead to a hypoplasia and loss of the Purkinje cells. Abnormal motor symptoms in TAF1-edited rats were associated with irregular cerebellar output caused by changes in the intrinsic activity of the Purkinje cells. Immunostaining revealed a reduction in the expression of the CaV3.1 T-type calcium channel. This animal model provides a powerful new tool for studies of neuronal dysfunction in conditions associated with TAF1 abnormalities and should prove useful for developing therapeutic strategies to treat TAF1 ID syndrome.

**Significance Statement:** Intellectual disability (ID) syndrome is an X-linked rare disorder caused by loss-of-function mutations in the *TAF1* gene. There is no animal model for understanding neuronal pathology and to facilitate development of new therapeutics for this X-linked intellectual disability syndrome. Novel animal models are valuable tools for understanding neuronal pathology and to facilitate development of new therapeutics for diseases. Here we developed a novel animal model for TAF1 ID syndrome in which the *TAF1* gene is deleted by CRISPR-Cas9 editing and lentiviral transduction. This animal model provides a powerful new tool for studies of neuronal dysfunction associated with TAF1 abnormalities and should prove useful for developing therapeutic strategies to treat TAF1 ID syndrome.

## Introduction

Mutations in TATA-box binding protein factor 1 (TAF1) protein are associated with a X-linked TAF1 intellectual disability (ID) syndrome (MIM: 300966) (2, 3). Occurring in males, TAF1 ID syndrome presents with abnormalities in global developmental (motor, cognitive, and speech), hypotonia, gait abnormalities, and cerebellar hypoplasia (2, 3). Exactly how mutations in TAF1 give rise to these neurological deficits remain unclear and there are no effect treatments for this disorder. Because TAF1 appears to serve as a scaffold on which TAF binding proteins and other TAFs interact in the assembly of transcription factor TFIID (4, 5), it is not known how the many functions of this protein complex are linked to the development of TAF1 ID syndrome Before therapeutic options for the treatment of TAF ID syndrome can be evaluated, an animal model for this disease needs to be developed. Thus, recapitulating major clinical features and disease progression reported in TAF1 ID patients would be beneficial.

We hypothesized that deletion of T*AF1* by CRISPR/Cas9 editing, into the maturing brain during the post-natal days (6–8) using a lentiviral vector might replicate TAF1 ID symptoms. This mode of administration allows the vector diffusion and infection to occur before maturation of axon myelination and gliogenesis – two processes that restrict free diffusion in the brain as well as infection of the maturing cerebellum. Similar approaches have been utilized to develop a rat model of synucleinopathy (9) and dystonia DYT1 (10). We found that in neonatal rat pups, removal of *TAF1* by CRISPR/Cas9 editing, results in defects in neonatal motor functions. Furthermore, the motor deficits were associated with loss of Purkinje cells in the cerebellum; the remaining Purkinje cells displayed abnormal firing frequencies. In addition, the motor defects persisted in *TAF1*-edited juvenile pups and were associated with morphologic abnormalities within the cerebellum.

## Results

### Design of CRISPR/Cas9 strategy for *TAF1* gene editing in the CNS

We designed a clustered regularly interspaced short palindromic repeats (CRISPR) associated protein-9 nuclease (Cas9) approach to induce a double stranded break in the promoter region of the *TAF1* gene (Fig. 1A), which would lead to reduced expression of TAF1 protein. The primary reasons for selecting this region of the gene is that patients with TAF1 variants present with loss of function abnormalities (2, 3). Lentiviral particles were generated and were injected intracerebroventricularly (ICV) into the brain of neonatal rats. Injection sites were verified with Evans blue dye (Fig. S1). Red fluorescent protein (RFP) staining indicated that cells within the cerebellum including Purkinje cells were transduced but not naïve animals (Fig. 1B i-iii). Using an antibody specific for TAF1, we detected efficient genome editing of the *TAF1* gene resulting in reduced TAF1 protein expression (Fig. 1C i-iii).

**Figure 1.**
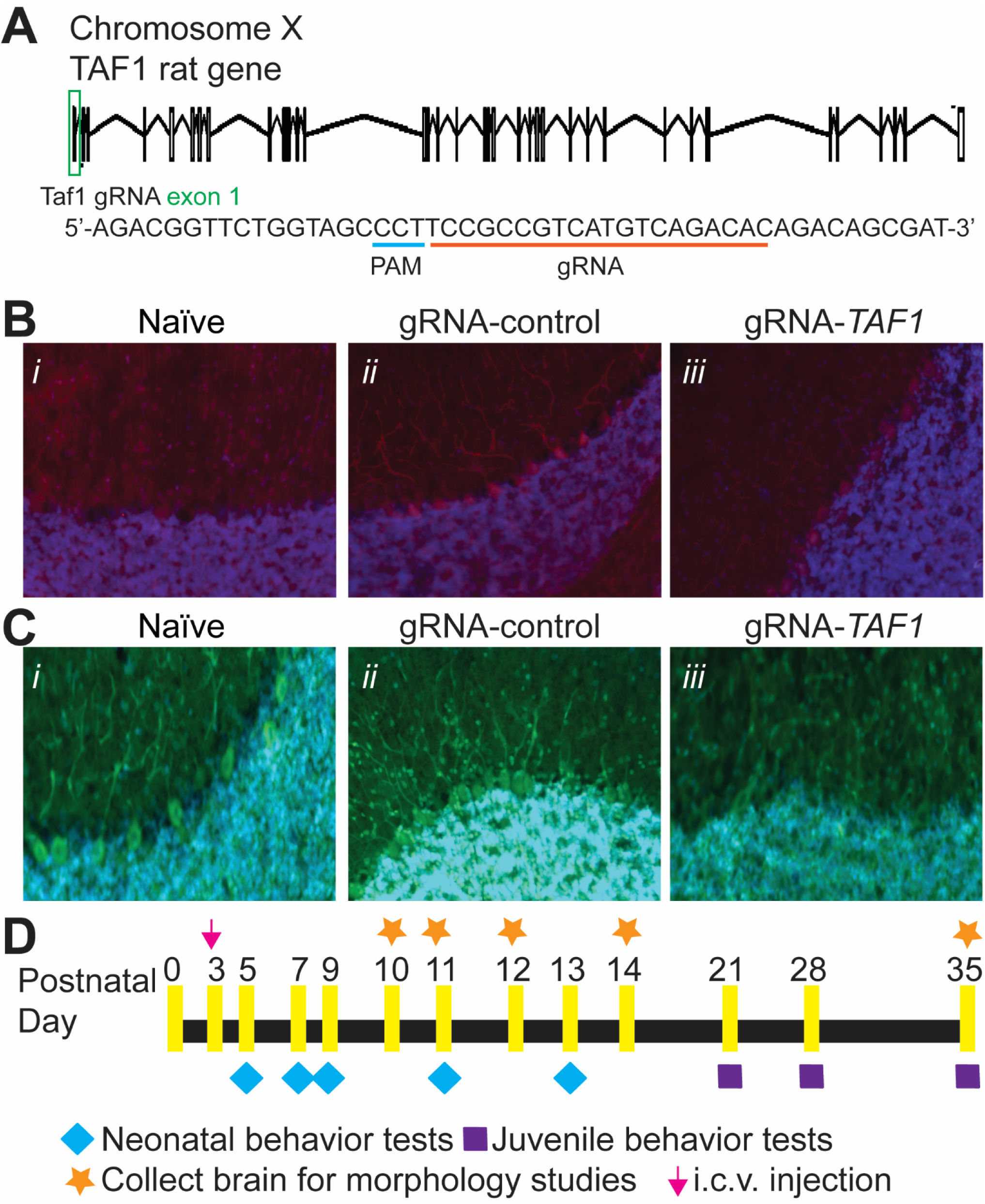
Validation of *TAF1* gene editing in cerebellar Purkinje cells and behavioral experimental design. (*A*) Schematic showing exon/intron organization of the *TAF1* gene in rat. To knockout TAF1, we designed a gRNA targeting exon 1. We used a guide RNA (gRNA) targeting a non-coding region in intron 1 of the *TAF1* gene as a control. Both gRNAs are based on the reverse strand sequence and are indicated on the figure by the red line oriented from 5’ to 3’. The protospacer adjacent motif (PAM sequence) is indicated by a blue line. The gRNA pairs with its DNA target followed by a 5’NGG sequence (PAM sequence). Cas9 catalyzes a double-stranded cleavage on the genomic DNA 3 bp before the PAM sequence. Nucleotide positions are indicated based on the DNA sequence on the *TAF1* gene. Representative immunohistochemistry images of cerebellar Purkinje cells stained with an antibody against red fluorescent (RFP) protein (*B*) or *TAF1* (*C*). Images were obtained from slices of rat pups uninjected (naïve) or injected (with gRNA-control or gRNA-TAF1) at 35 days post-injection. Reduced TAF1 expression was found in the *TAF1*-edited rat pups compared to control group rat pups. These experiments were performed with 4 male animals per each experimental condition in a double blinded manner. Scale bar = 200 μm. (*D*) Experimental design for neonatal and juvenile motor tests and histopathology studies.

### CRISPR/Cas9-mediated TAF1 deletion cause TAF1 ID-like motor deficits

Motor dysfunction is a characteristic of TAF 1D syndrome (2, 3). Therefore, we performed a battery of neonatal motor tests to ascertain possible motor deficits between TAF-edited rat pups and control animals (Fig. 1D). Negative geotaxis refers to the orienting response and movement expressed in opposition to cues of a gravitational vector (11). We observed that *TAF1*-edited pups displayed increased latency time in correcting their position relative to controls (Fig. 2A). The righting reflex evaluates the motor ability for a rat pup to flip onto its feet from a supine position (11). Similar to the negative geotaxis test, *TAF1*-edited pups showed an increased latency time to “right” their position (Fig. 2B). The tail suspension test is commonly used to measure stress and can also be used to assess motor alterations (i.e. motor coordination). In the tail suspension, the naïve and gRNA-control animals held their hind limbs wide apart from PD5 to PD13, whereas the *TAF1*-edited rats exhibited abnormal hind-limb clasping after PD7 (Fig. 2C). The hind-limb suspension test assesses the proximal hind limb muscle strength, weakness, and fatigue in the animal. The hind-limb suspension test also evaluates general neuromuscular function, body muscle strength, and posture. It is designed specifically for neonates and can be used on animals PD 2-14 (12). *TAF1*-edited rat pups showed hind-limb weakness as demonstrated by a marked decrease in hanging score compared to control animals (i.e. gRNA-control and naïve rat pups at the same age) (Fig. 2D). Crawling is a behavior developed early in rodent pups between postnatal day (PD) 0-5, at which point pups start to transition to walking from 5-10 days old (11). Thus, the ambulation test takes advantage of this transitional time course. We noted significant differences in ambulation, at PD 14, between the *TAF1*-edited rat pups compared to gRNA-control and untreated (naïve) rat pups (Fig. 2E). Collectively, these data indicate motor dysfunction in CRISPR/Cas9-*TAF1*-edited treated rat pups.

**Figure 2.**
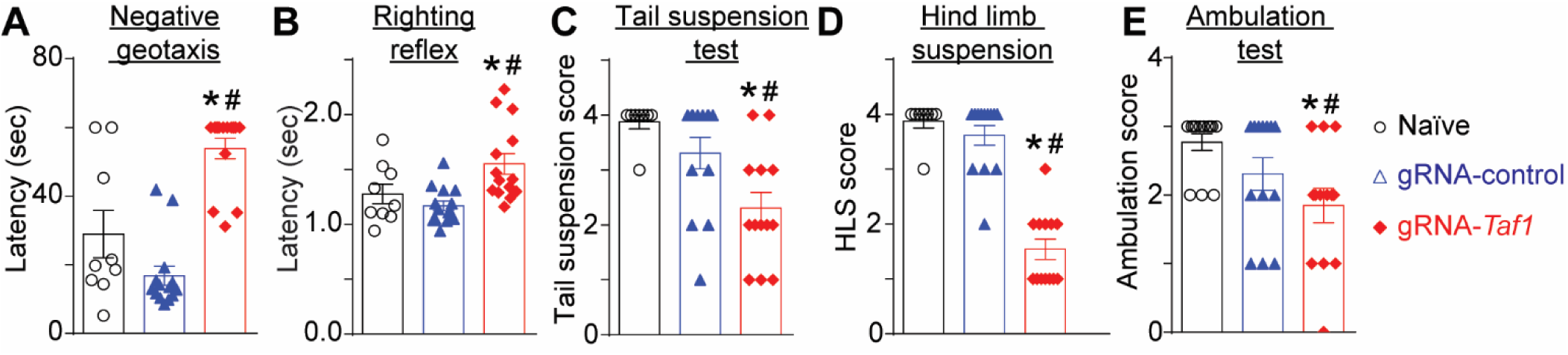
Behavioral assessment of neonatal motor functions. The *TAF1*-edited rats showed increased latency time in negative geotaxis (*A*) and righting reflex test (*B*). Similarly, the TAF1-edited rat pups demonstrated decreased scores in the tail suspension (*C*), hind limb suspension (*D*), and ambulation (*E*) tests. Data are shown as mean ± S.E.M., n = 14 per condition. *p<0.05 versus; naïve, #p<0.05 versus gRNA-control (ANOVA followed by Tukey’s test). The experiments were conducted in a blinded fashion.

We continued our behavioral assessment past the weanling stage. We performed the beam walking test and Open field test. The beam walking test can evaluate motor coordination and balance (13). The beam crossing time (Fig. 3A) and number of foot slips errors were higher in the *TAF1*-edited animals at PD35 compared to controls (Fig. 3B). In the open field test, *TAF1*-edited animals displayed a higher level of anxiety as inferred from increased grooming frequency (Fig. 3C). Thus, *TAF1*-editing produces behavioral defects that appear to persist into the juvenile animals.

**Figure 3.**
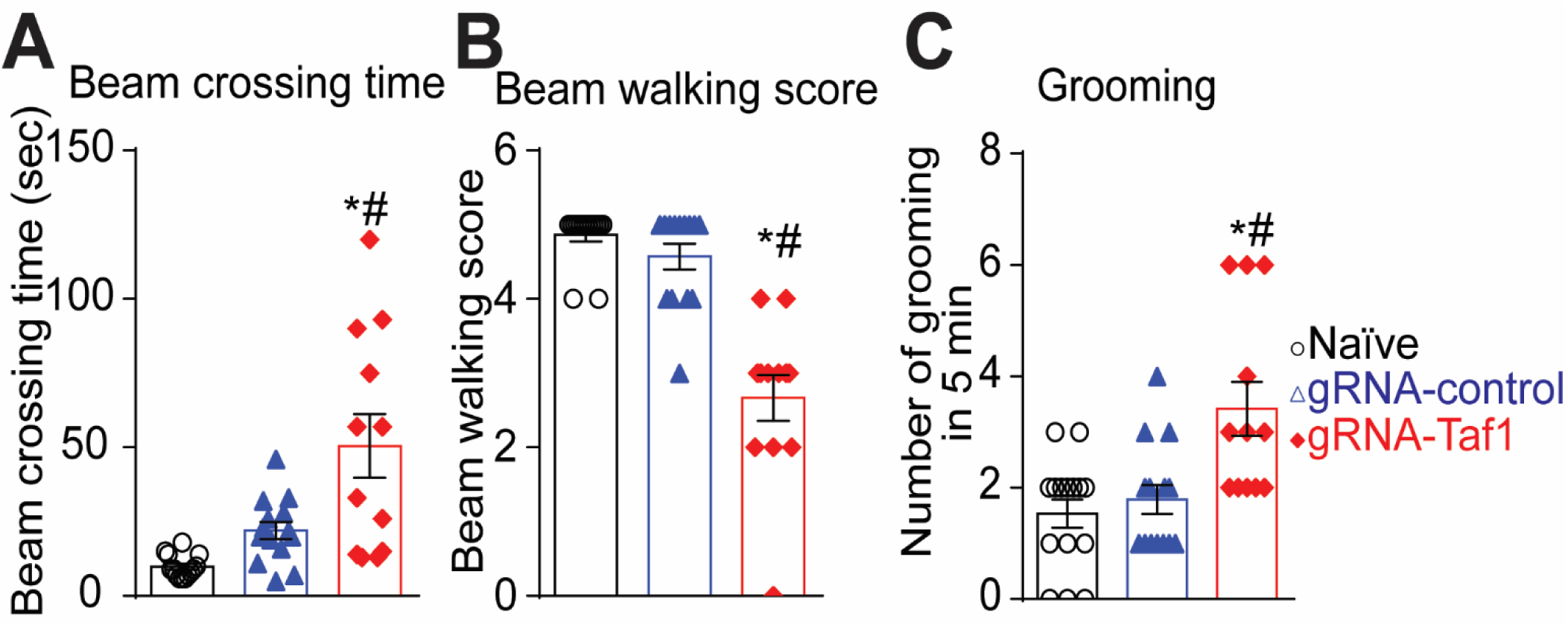
Behavioral assessment of juvenile motor functions. The *TAF1*-edited animals showed behavior deficits compared to naïve and gRNA-control group animals. The *TAF1*-edited showed increased beam waking time (*A*) and had a decreased beaming score (*B*) in beaming walking tests relative to naïve and gRNA-control rats. (*C*) Grooming was increased in the *TAF1*-edited rats compared to the other groups. Data are shown as mean ± S.E.M., n = 5-8 per experimental condition. *p<0.05 versus; naïve, #p<0.05 versus gRNA-control (ANOVA followed by Dunnett’s test). The experiments were conducted in a blinded fashion.

### TAF1 deletion leads to cerebellar abnormalities

To examine a possible cellular basis for the behavioral defects, next we evaluated the morphology of the Purkinje cells by Hematoxylin and eosin (H&E) and Nissl staining. Our histopathology analysis showed that the architecture of the Purkinje cell layer was abnormal in *TAF1*-edited animals. In addition, the histopathology showed Purkinje cells were markedly hypoplastic in TAF1 treated animals relative to controls (Fig. 4A and 4B i-iii).

**Figure 4.**
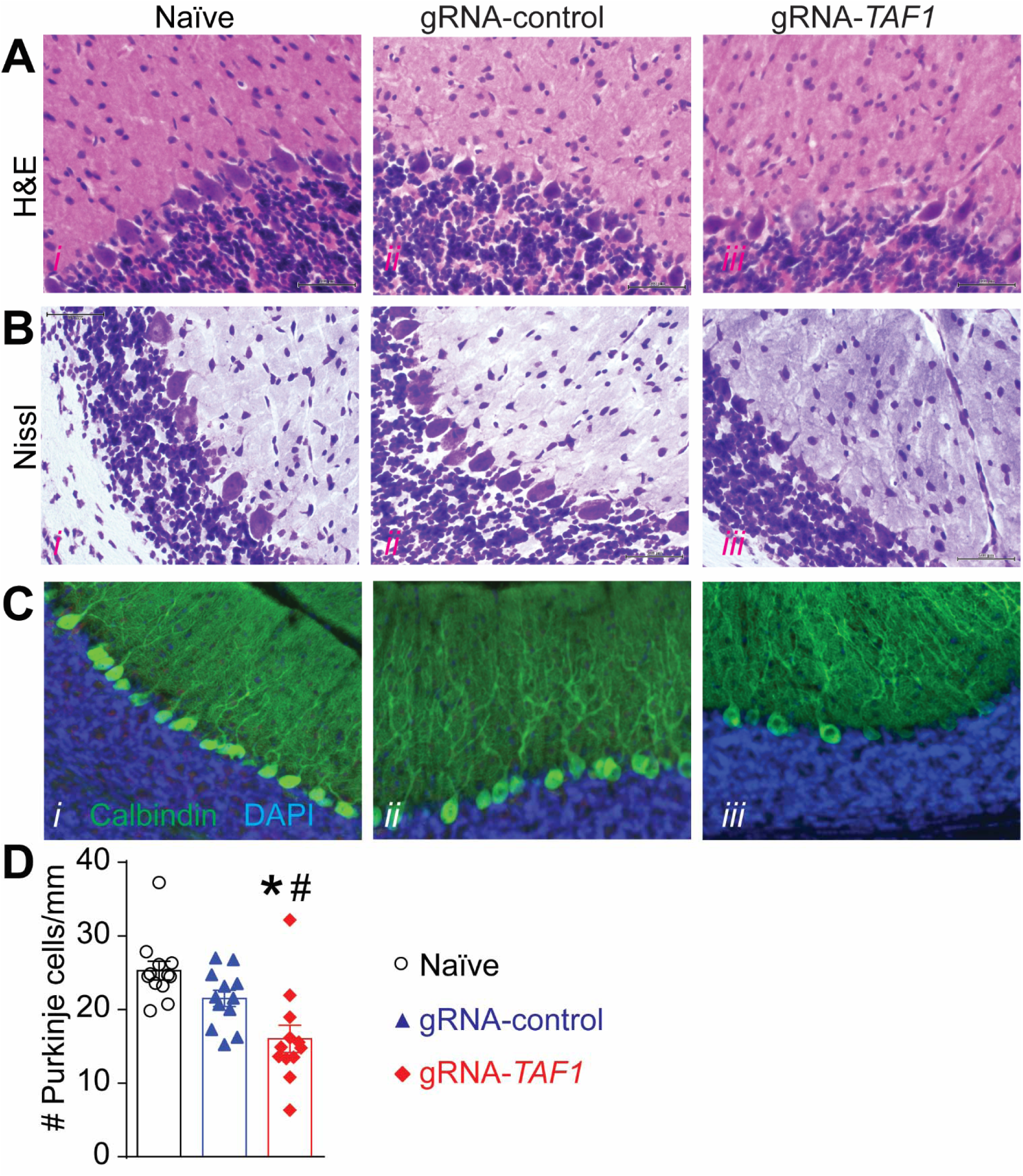
*TAF1* gene editing results in loss of cerebellar Purkinje cells. The morphology of the Purkinje cell layer was evaluated by Haemotoxylin and Eosin (*A*) and Nissl (*B*) Staining. Both H&E and Nissl staining showed abnormal architecture of the Purkinje cells as well as hypoplastic Purkinje cells. Scale bars: 500 μm. (C) Expression of calbindin was decreased in *TAF1*-edited animals as compared to naïve and CRISPR-control groups. Note also a decrease in the number of Calbindin positive Purkinje cells in TAF1-edited animals compared to control animals. (*D*) Summary of the number of Purkinje cells per linear density in each of the experimental conditions. Data are shown as mean ± S.E.M., n = 12 fields per animal, 4 animals per experimental condition. *p<0.05 versus; naïve, #p<0.05 versus gRNA-control (ANOVA followed by Tukey’s test). The experiments were conducted in a blinded fashion.

Purkinje cells are characteristically Calbindin positive and the protein is abundant in the cell body axon and dendritic tree of the naïve and gRNA-control animals, however some of the Purkinje cells in the TAF1-edited animals did not stain as positive as the control group animals. (Fig. 4C i-iii). We also observed a decrease in the number of Calbindin positive Purkinje cell in TAF1-edited animals compared to control animals (*Fig. D*). We conclude that organization of the Purkinje cell layer and morphology of Purkinje cell are altered during postnatal development in *TAF1*-edited rats.

### CRISPR/Cas9-mediated TAF1 deletion leads to Purkinje cells excitatory neurotransmission abnormalities

To examine a possible functional basis for the behavioral defects, we next performed synaptic recordings of neurons in the Purkinje cell layer region of cerebellar slices (Fig. 5A) using whole-cell patch clamp, specifically measuring spontaneous excitatory postsynaptic currents (sEPSCs) (Fig. 5B). There was no change in amplitude of sEPSCs between three groups (Fig. 5B-D). However, the frequency of sEPSCs was significantly decreased with *TAF1*-editing, compared with naïve or gRNA-control editing, respectively (Fig. 5F) (1.61 ± 0.40 Hz vs 4.67 ± 0.54 Hz, *P<0.05 vs Naïve; 1.61 ± 0.40 Hz vs 4.30 ± 1.00 Hz, #P<0.05 vs gRNA-control). These results confirm a presynaptic effect on glutamatergic transmission of cerebellar Purkinje cells with *TAF1*-editing.

**Figure 5.**
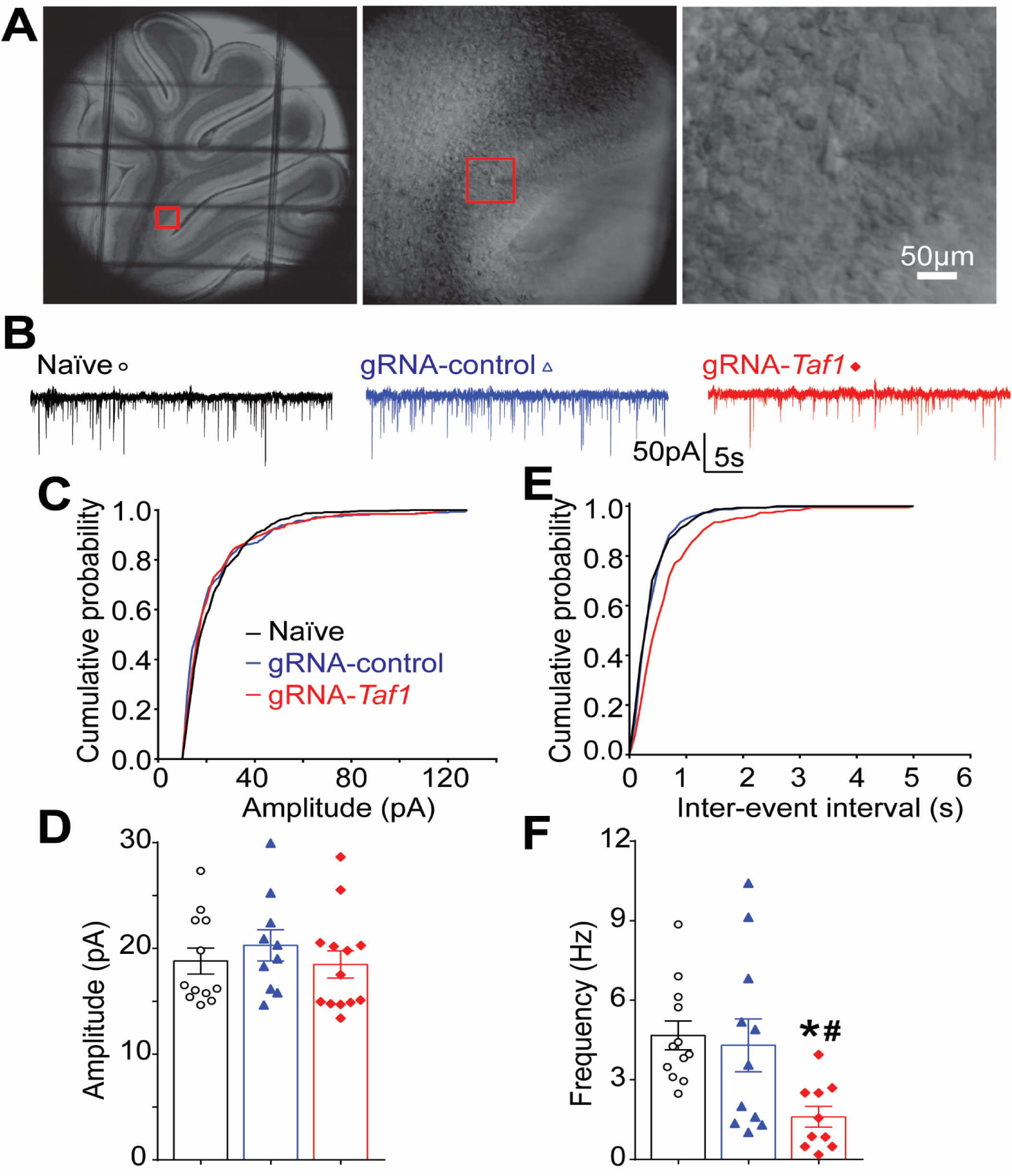
*TAF1* gene editing leads to a decrease in frequency of spontaneous excitatory post synaptic current in cerebellar Purkinje cells. (*A*) Photomicrograph of cerebellar slice preparation with a progressive zoom with the rightmost panel showing positioning of the recording electrode to this region. (*B*) Representative recording traces of cells from the indicated groups. The cumulative probability of amplitude (*C*) and inter-event interval (*E*). Summary of amplitudes (*D*) and frequencies (*F*) of sEPSCs for the indicated groups are shown. Data are shown as mean ± S.E.M., n = 12 Purkinje cells from at least 2 animals per experimental condition. *p<0.05 versus; naïve, #p<0.05 versus gRNA-control (ANOVA followed by Tukey’s test). The experiments were conducted in a blinded fashion.

### CRISPR/Cas9-mediated *TAF1* deletion is associated with reduced CaV3.1 protein expression

T-type calcium (Ca^2+^) channels play a key role in regulating membrane excitability in the brain (14–16). In the cerebellum the T-Type channel, CaV3.1, is expressed in Purkinje cells and appears to be involved in the entry of Ca^2+^ into presynaptic terminals (14, 17, 18). Because we previously reported altered Ca^2+^ entry and reduced protein expression of Cav3.1 in *TAF1* depleted SH-SY5Y cells (2), we investigated if Cav3.1 expression was altered in vivo. We observed comparable levels of expression of CaV3.1 in Purkinje cell slices from both naïve animals and gRNA-control injected animals, whereas CaV3.1 expression was markedly reduced in *TAF1*-edited animals at post-natal day 14 (data not shown) and at post-natal day 35 (Fig. 6A).

**Figure 6.**
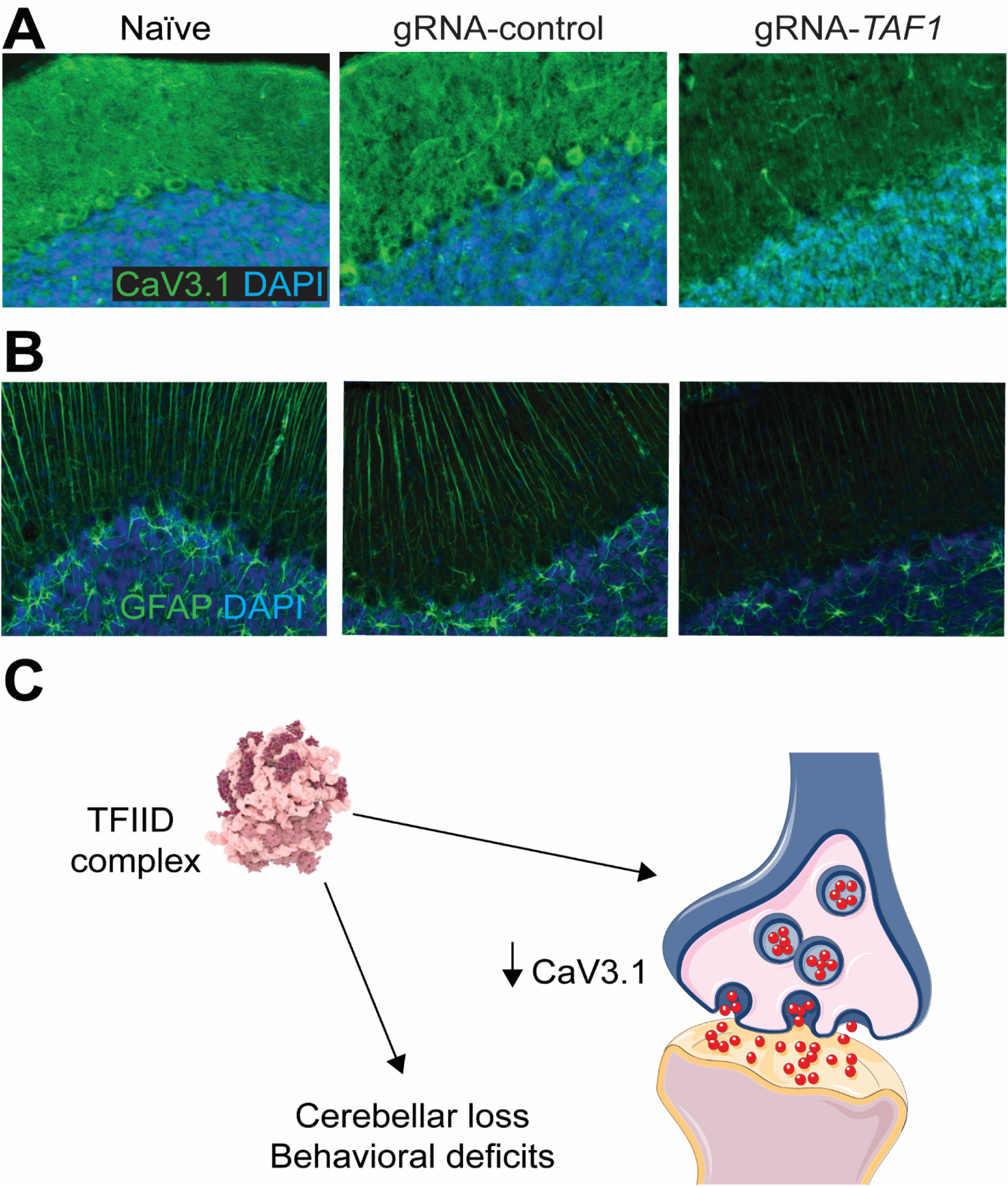
*TAF1* gene editing leads to a downregulation of the CaV3.1 voltage-gated calcium channel. (*A*) Representative photomicrographs of cerebellar slices from animals injected with (vehicle) or control or TAF1 gRNAs stained with an antibody against calbindin. Nuclei stained with DAPI. (*B*) Proposed summary of deletion of the *TAF1* gene causing cerebellar hypoplasia (similar to that seen in humans (2, 3)) in rats along with a reduction in CaV3.1 calcium channels. CRISPR/Cas9-mediated TAF1 deletion leads to Purkinje cells excitatory neurotransmission abnormalities which lead to behavioral motor abnormalities.

## Discussion

Mutations in TAF1, which is a component of the TFIID transcription factor complex, are implicated in an X-linked genetic syndrome that afflicts males (3). Some of the clinical features associated with TAF1 ID syndrome include: delayed gross motor development, hypoplasia of the cerebellar vermis, and hypotonia (2, 3). By deleting the *TAF1* gene in postnatal rats, we generated an animal model that replicates the clinical features of TAF1 ID syndrome. We observed motor deficits in both young rats and juveniles, morphological changes and a loss in Purkinje cells in the *TAF1*-edited animals. As in humans, we noted hypoplasia of the cerebellum, a pathological hallmark of dysfunction in TAF1 ID syndrome. Our model underscores the relevance of the cerebellum to this neurodegenerative disease and offers a powerful platform for examination of the cellular and molecular pathways underpinning TAF1 ID syndrome.

Our model of TAF1 ID allowed us to explore how cerebellar dysfunction contributes to disease. Two different pathways, parallel fibers (PF) from granule cells (GC) and climbing fibers (CF) from the cells in the inferior olive (I.O), project excitatory synaptic input to Purkinje cells (PC)(19). PCs integrate these excitatory inputs and then send inhibitory GABAergic projections to the deep cerebellar nuclei (DCN) (20). We found that, in Purkinje cells, the frequency of sEPSCs was significantly decreased with *TAF1*-editing, compared with naïve or control CRISPR respectively. These results are suggestive of presynaptic suppression of *TAF1*-editing on glutamate transmission of cerebellar Purkinje cells, which might lead to disinhibition of PCs and the alterations in ongoing motor activity and fine motor control. We found that the organization of Purkinje cell layer and morphology of Purkinje cell are altered during postnatal development in *TAF1*-edited rats, we infer that the decrease of frequency of sEPSCs might result from the collapse of the cerebellar circuit.

T-type calcium channels are low voltage-activated calcium channels that transiently open to evoke tiny Ca^2+^ currents (reviewed in (15)). T-type calcium channels regulate calcium influx from the extracellular region by opening the calcium channel (21), or activating calcium-induced calcium release from the internal calcium source (22, 23). These results suggest a critical role for T-type calcium channels in regulating intracellular calcium homeostasis and maintaining cellular function (21, 24, 25). CaV3.1 T-type channels are abundant at the cerebellar synapse between parallel fibers and Purkinje cells where they contribute to synaptic depolarization (26). Consistent with our previous in vitro findings (2), we observed a decrease in CaV3.1 expression in Purkinje cells in the present study. Because CaV3.1 appears to be required for neuroprogenitor cell viability and the primary pathway by which Ca2+ enters into cerebellar neurons (26, 27), our results suggests that dyregulated calcium signaling, due to decreased CaV3.1 protein levels caused by *TAF1* gene knockout, could be important in the pathogenesis of TAF1 ID syndrome.

In order for a gene to be transcribed, a transcriptional preinitiation complex (PIC) needs to assemble at its promoter. The first complex to bind the promoter is the general transcription factor, TFIID, consisting of the TATA-binding protein (TBP) and 13 different TBP-associated factors (TAFs) (5, 28). Mutations in several other human TFIID subunit coding genes (i.e. *TBP*, *TAF2*, *TAF6*, TAF8, and *TAF13*) have been implicated in human diseases including many with neurological outcomes and intellectual disability (29–33). The outcome of these mutations on TAFs has not been assessed in detail, due in part to the lack of the development of animal models for these disorders of transcription regulation as disruption of TFIID subunits can be embryonic lethal (32, 34). Our facile approach for assessing TAF1 function *in vivo*, using CRISPR/Cas9-based gene editing coupled lentiviral transduction, could be applied to these disorders of transcription regulation and provide insight into underlying mechanisms of disease. Although our work did not achieve germline deletion of TAF1, the approach used here provides an instructive example of how spatially and temporally restricted deletion may be sufficient to mimic hallmarks of these neurological disorders.

In conclusion, the present study shows that targeted knockdown of *TAF1* can be achieved in the brain of rats with postnatal delivery of a CRISPR-Cas9 lentiviral vector. The current results demonstrate that this approach led to motor defects and morphological abnormalities within the cerebellum reminiscent of the human neurological disease-TAF1 ID syndrome. This model therefore provides an interesting opportunity for further studies investigating the effects of aberrant TAF1 on neuronal dysfunction and paves the way for testing of possible therapeutic strategies to treat TAF1 ID syndrome.

## Materials and Methods

Detailed descriptions of methods used, and any associated references are available in SI Materials and Methods. Briefly, all biochemical, electrophysiology and behavior experiments were performed in a blinded fashion according to established protocols. Animal protocols were approved by the Institutional Animal Care and Use Committee of the College of Medicine at the University of Arizona and conducted in accordance with the Guide for Care and Use of Laboratory Animals published by the National Institutes of Health.

## Supporting information

Supplementary_methods-Taf1

## Author Contributions

MAN and JU designed research; AA, JU, AM and SNB performed research; JU, MAN, and RK analyzed data; and MAN, JU, and RK wrote the paper.

## Acknowledgements

We would like to thank Drs. Todd Vanderah and Tally Largent-Milnes for the ICV injection facilities and behavior analysis. We would also like to thank Robert Hershoff for his help with the digital images. We also acknowledge the funding and support of the Senner Endowment for Precision Health, University of Arizona Health Sciences. This work was also supported by grants from the National Natural Science Foundation of China (81603088) to J.Y., and R01DA042852 from the National Institute on Drug Abuse to R.K.); and a Neurofibromatosis New Investigator Award from the Department of Defense Congressionally Directed Military Medical Research and Development Program (NF1000099) to R.K.

## Conflict of interest

The authors have no conflict of interests to disclose.

---

**Fig. S1.**
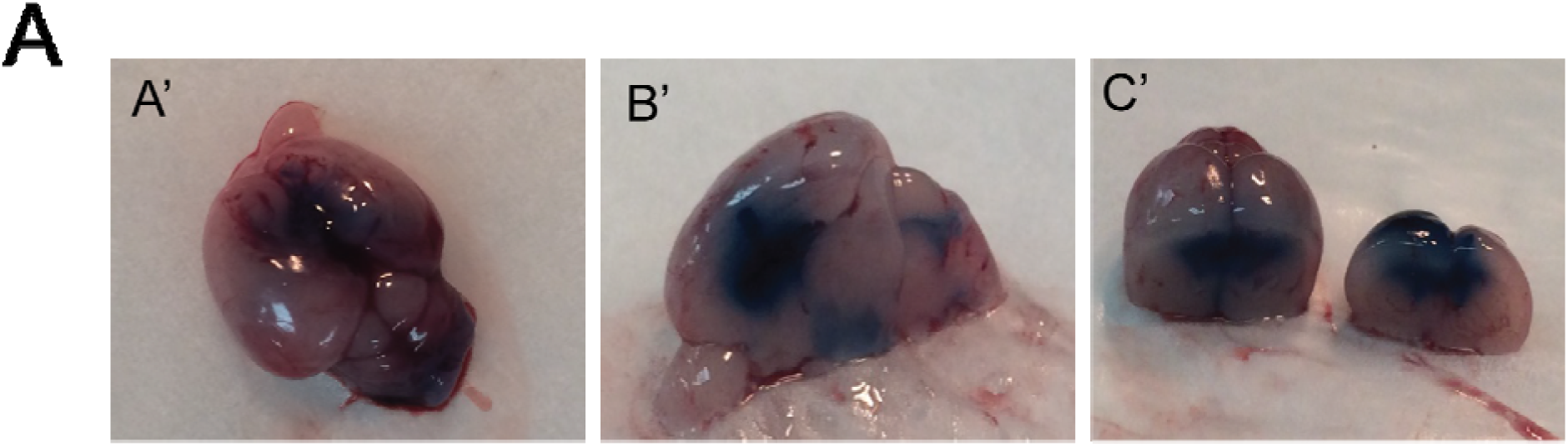
Evan’s blue migration and Nissl staining in cerebellum. (A) Evan’s blue migration from the ICV injection site to the cerebellum of PD 3 rat pups (A’) whole brain; (B’) sagittal section of brain; (C’) coronal section of brain. (B) Nissl staining (40x magnification) of cerebellar tissue from Naïve, CRISPR-control and CRISPR-TAF1 rats. Normal morphology of the Purkinje cells in Naïve and CRISPR-co trol rats. Whereas the morphology of the Purkinje cell in the CRISPR-TAF1 treated rats was abnormal and characterized by hypo plastic PCs and loss of architecture of the PCs layer.

