## Supplementary_methods-Taf1 for "*TAF1*-gene editing impairs Purkinje cell morphology and function"

**Supplentary Figure and Methods**

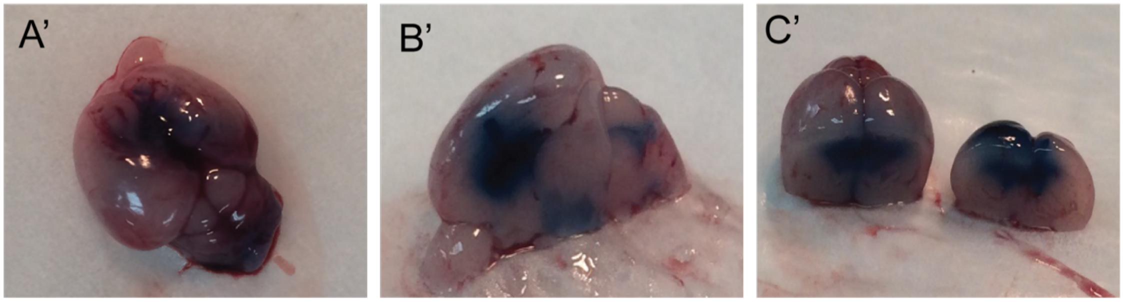


**Supplementary Figure 1.** Representative Evan's blue staining showing migration from the ICV injection site to the cerebellum of PD 3 rat pups and the Nissl staining (40x magnification) of naïve and experimental rats PCs. Rats were injected with vehicle (*A'*), gRNA-control (*B'*), or gRNA-TAF1 (*C'*).

### Materials and Methods

#### Animals

Pathogen-free, normal E18 pregnant Sprague–Dawley rats (Envigo Laboratories) were housed 1 per cage in temperature- (23 ± 3 °C) and light (12-h light/12-h dark cycle; lights on 07:00–19:00)-controlled rooms with standard rodent chow and water available ad libitum. The neonates are designated as P 0 on the day of birth, the litter size range included in current study was 9-14 pups. For all experiments, the experimenter was blinded to the treatments.

#### gRNA strategy for *TAF1* gene targeting.

Our strategy to truncate neurofibromin focused on targeting exon 1 of the *TAF1* gene using a guide RNA (gRNA) as described previously (1-4). We targeted this exon to ensure total removal of the Taf1 protein. Using this approach, we expect minimal to none off-target activity of the Cas9 enzyme as we and others verified before (1, 5). The gRNA sequence (GTGTCTGACATGACGGCGGA, quality score 94) was inserted into the restriction site of the pL-CRISPR.EFS.tRFP lentiplasmid (Cat#57819, Addgene, Cambridge, MA) (6) a plasmid that allows for simultaneous expression of (i) the Cas9 enzyme; (ii) the gRNA; and (iii) a red fluorescent protein (tRFP) – to control for transduction efficiency. All plasmids were verified by Sanger sequencing (Eurofins, Louisville, KY). Lentiplasmids were packaged in lentiviruses by Viracore (USCF) at titers routinely above $10^7$ infectious particles per ml.

#### Intracerebroventricular injections

All animal procedures were approved by the University of Arizona Institutional Animal Care and Use Committee and are in accordance with the NIH Guide for Care and Use of Laboratory Animals. Bilateral intracerebroventricular (ICV) injections were performed as previously described in Sprague-Dawley rat pups on postnatal day 3 (7) (8). Briefly, Newborn SD rat puts were anesthetized by isoflurane. A 10 µl syringe (Hamilton Gastight Syringe, #1701) was used to pierce the skull (coordinates from bregma: -0.6 mm posterior, ± 1.75 mm lateral/medial, and -2.5 mm ventral), and 2.5 µl of CRISPR lentivirus (gRNA-control or gNRA-*TAF1*) was injected into each cerebral ventricle without opening the scalp. Neonatal rat pups were kept with parent until weaned.

Rats were sacrificed at set time points as follow: 14 days and 35 day of postnatal day. Of note, 14 day and 35 day groups were behaviorally assessed before being euthanized for electrophysiological and histological and protein expression analysis.

**Cerebellar Slice Preparation**

10- to 14-day SD rat pups were used in the electrophysiological experiment as before (9). All animal procedures were carried out in accordance with the regulations of the Institutional Animal Care and Use Committee of University of Arizona (Tucson, USA). The animals were decapitated after being deeply anesthetized with isoflurane, and their cerebellums were rapidly removed and placed in an ice-cold dissection ACSF containing (in mM) 220 sucrose, 2.5 KCl, 1.25 Na2HPO4, 3.5 MgCl2, 0.5 CaCl2, 25 NaHCO3, and 20 D-glucose (with pH at 7.4 and osmolarity at 310 mOsm), bubbled with 95% O2 and 5% CO2. Parasagittal slices (320 μm thick) were cut using a VT 1200S vibratome (Leica, Germany). Slices were then incubated for at least 1 hour at 34°C in an oxygenated recording solution containing (in millimolar): 125 NaCl, 2.5 KCl, 2 CaCl2, 1 MgCl2, 1.25 NaH2PO4, 26 NaHCO3, 25 D-glucose, with pH at 7.4 and osmolarity at 320 mOsm. The slices were then positioned in a recording chamber and continuously perfused with oxygenated recording solution at a rate of 1.5 to 3 mL/min before electrophysiological recordings at RT.

**Whole Cell Patch-clamp Recording After incubation**

Recordings were made from PCs in lobules IV-VI, which were visually identified based on their location using infrared differential interference contrast video microscopy on an upright microscope (FN1; Nikon, Tokyo, Japan) equipped with a 3 40/0.80 water-immersion objective and a charge-coupled device camera. The pipettes were prepared by pulling glass capillaries on a model p-97 microelectrode puller (Sutter Instrument, Novato, USA). Patch pipettes had resistances of 3-5 MΩ. The internal solution was a K-based solution containing (in mM): The pipette solution contained the following (in millimolar): 120 potassium gluconate, 20 KCl, 2 MgCl 2, 2 Na 2 - ATP, 0.5 Na-GTP, 20 HEPES, 0.5 EGTA, with pH at 7.28 (with potassium hydroxide [KOH]) and osmolarity at 310 mOsm.

The whole-cell configuration was obtained in voltage-clamp mode. The membrane potential was held at -70 mV using PATCHMASTER software in combination with a patch clamp amplifier (EPC10; HEKA Elektronik, Lambrecht, Germany). To record spontaneous excitatory postsynaptic currents (sEPSCs), bicuculline methiodide

(10 m M) was added to the recording solution to block γ-aminobutyric acid-activated currents. Hyperpolarizing step pulses (5 mV in intensity, 50 milliseconds in duration) were periodically delivered to monitor the access resistance (15-25 MΩ), and recordings were discontinued if the access resistance changed by more than 20%. For each PC, sEPSCs were recorded for a total duration of 2 minutes. Currents were filtered at 3 kHz and digitized at 5 kHz. Data were further analyzed by the Mini-Analysis (Synatosoft Inc, NJ) and Clampfit 10.7 Program. The amplitude and frequency of sEPSCs were compared between neurons from three groups.

**Neonatal Motor Test**

At each testing day, the dams and the neonates were brought to the experimental room at the same time and left undisturbed for at least 1 h prior to testing. Then, the neonatal motor tests were performed as indicated below.

**Righting reflex test**

This test evaluates the body's tendency to regain its former body position after being displaced. The pups were placed on their backs on a flat surface (10). Their success or failure in repositioning themselves dorsal side up and the latency to falling off the platform were assessed over a 30 s period. For mice that failed to fall off, the time was recorded as 30 s. The measures were taken once every second day from PD5 to PD13.

**Negative Geotaxis**

This test evaluates motor coordination and vestibular sensitivity. Negative geotaxis was tested by placing the neonate on an inclined grid (at approximately 35° inclination) with the head facing downward (11). The grid had a rough surface to provide the neonate with a reasonable grip. We assessed the percentage of pups that successfully completed the test at each postnatal day and their latency to complete the geotaxis test in seconds (maximum 60 s). For those mice that failed to reorient themselves upward head, the time was recorded as 60 s. The measures were taken once every second day from PD5 to PD13.

**Ambulation Test**

Crawling is a behavior developed early in the mouse pup between PND 0 - 5, at which point mice begin to transition to walking between 5 - 10 days old (12). Pups were placed in a clear enclosure where they are visible from the top as well as from the side. The ambulation was scored for 3 min using the following scale: 0 = no movement, 1 = crawling with asymmetric limb movement, 2 = slow crawling but symmetric limb movement, and 3 = fast crawling/walking.

**Hind-limb Suspension Test**

This test was used to assess the general neuromuscular function and posture. We used the method of EL-Khodor to evaluate the hind-limb muscle strength, weakness, and fatigue in rat neonates (13). In each trial, the pups were placed head-down, hanging by their hind limbs in a plastic 50 ml centrifuge tube with a cotton ball cushion at the bottom to protect the animal's head upon its fall. The test was assayed over a 40-s period. The hind-limb score that assessed the positioning of the legs. The posture adopted by the pups was scored according to the following criteria: a score of '4' indicated normal hind-limb separation; a score of '3' indicated that the hind limbs were closer together than normal, without touching; a score of '2' specified closer proximity of the hind limbs to each other, often touching; and a score of '1' represented touching or clasping of the hind limbs. A score of 1 was given to a pup that failed to maintain its grasp of the tube. The score is an overall evaluation of the hind-limb position during the first 10 s of hanging onto the lip of the tube. The measures were taken once every second day, from PD5 to PD13.

**Tail Suspension Test**

The pups were suspended by their tails for 5-s and their hind-limb postures were scored 1–4, as defined in (14). The measurements were taken every other day, from PD5 to PD13.

**Juvenile Motor Test**

After weaning the rat pups were separated from the dams and maintained 4 per cage. At each testing day, rats were brought to the experimental room at the same time and left undisturbed for at least 1 h prior to testing.

**Open Field Test**

The apparatus (W100 cm×D100 cm×H40 cm) is made of wood and resin. The floor of this chamber was divided into 25 cm (5 × 5) squares. The rat at indicated ages were placed into one corner of an open field chamber and their behavior was observed for 5 min as before (9). The number of episodes for grooming (i.e. consisting of licking the fur), washing the face or scratching behaviors were quantified (15).

**Beam Walking Test**

Animals were allowed to walk on a narrow flat stationary wooden beam (L100 cm×W2 cm) placed at a height of 100 cm from the floor to reach an enclosed escape platform. The time taken to cross the beam from one end to the other and beam walking score were also given as described previously (15).

**Hematoxylin & Eosin Staining**

Sections of cerebellum (20-µm thick) were prepared and then stained with hematoxylin and eosin (H&E) dye, and mounted in neutral deparaffinated xylene medium for microscopic observations.

**Nissl Staining**

Sections of cerebellum (20 µm thick) were prepared and then stained with Cresyl violet dye, and mounted in neutral deparaffinated xylene medium for microscopic observations.

**Immunohistochemical staining and counting of Purkinje cells**

Rats were were perfused with 4% formaldehyde in PBS at PD 14 day and PD 35 day of age, and the brains were extracted and post-fixed for 8 h at 4°C. Cerebellar tissues were prepared and cut sagittally at 20 µm using a cryostat (Microm HM 505 E). After rinsing the sections in Phosphate buffer saline (PBS) for 5 min, the sections were incubated with a 0.1% H2O2 solution in PBS for 5 min, rinsed in PBS, and incubated for 30 min with 0.4% Triton X-100, rinsed in PBS and block with 8% goat serum, and 1% Triton X-100 in PBS. After blocking, the sections were incubated for overnight with anti-calbindin D-28 K polyclonal antibody (1:250, Millipore), anti-tubulin (1:250, Millipore), anti-TAF1 (1:250, Millipore), anti-tRFP (1:250, Evrogen) or anti-CAV3.1 (Alomone labs) as indicated, diluted in 4% goat serum in PBS. The sections were washed in 1% goat serum in PBS, incubated with secondary antibody (anti-rabbit Alexa fluora 488 & anti-mouse Alexa fluora 594) for 2 h, washed, and incubated with DAPI for 5 mins. Sections were then washed and further air dried, and cover slipped with Richard Allan Scientific Mountant Medium (Thermo). All procedures were performed at room temperature. Calbindin positive Purkinje cells were observed under a fluorescence microscope (LSM510, Carl Zeiss) using a 20 objective. Nucleated Purkinje cells in the anterior lobe (lobule I-III) of the cerebellar cortex in every fifth sections were counted using Icy software, and Purkinje cell linear density was calculated by dividing the number of cells by the linear distance of the Purkinje cell layer per section.

**Statistics**

Data were analyzed with the software Prism 6 (GraphPad). Group differences were calculated using ANOVA with Tukey's or Dunnett's multiple comparison tests as indicated in the figures. P values smaller than 0.05 were considered significant. Error bars in the graphs represent mean ± SEM.
